# A Guanine-quadruplex located on the negative strand of the hepatitis C virus facilitates efficient genomic RNA synthesis

**DOI:** 10.1101/2025.05.14.654041

**Authors:** Simmone D’souza, Scott Tersteeg, Maulik D. Badmalia, Annie Chen, Milan S. Heck, Guido van Marle, Jennifer A Corcoran, Carla S. Coffin, Trushar R. Patel

**Author notes:** (S.D.); (S.T.); (M.D.B.); (A.C.); (M.S.H.); (G.v.M.); (J.A.C.); Carla S. Coffin (C.S.C); (T.R.P.). Address correspondence to Carla S. Coffin (C.S.C) and (T.R.P.). Maulik D. Badmalia, University of Alberta, Edmonton, Alberta, Canada; Milan S. Heck, McGill University, Montreal, QC.

## Abstract

Guanine-rich nucleic acids can form a unique secondary structure called Guanine-quadruplexes (G4s). These G4s are conserved across all domains of life and in viruses, where they can play important regulatory roles in the viral lifecycle. In flaviviruses, the (-) strand 3’ untranslated region (UTR) is essential for initiating genomic RNA synthesis. Within the (-) strand 3’ UTR of hepatitis C virus (HCV), a highly conserved G4 is located within stem-loop IIy’ (SLIIy’), spanning nucleotides 110-131—a region that is essential for efficient replication. Using bioinformatics, we demonstrate that nucleotides 110-131 are highly conserved across HCV genotypes 1-7, and this region within SLIIy’ can likely adopt a more energetically favorable G4 conformation rather than the predicted hairpin. From biophysical and cell biology experiments, we show that stabilizing the predicted hairpin structure in SLIIy’ disrupts RNA synthesis, while successive guanine-to-adenine mutations within the G4 sequence impair G4 formation and hinder replication. Combining our findings with prior studies, we propose that the SLIIy’ G4 plays a crucial role in recruiting NS3 helicase, which is necessary for unwinding RNA structures and facilitating access for NS5B polymerase. The G4 may serve as a regulatory switch that modulates helicase binding and replication efficiency. We propose that in the absence of a stable G4, NS3 recruitment or unwinding is impaired, leading to inefficient RNA unwinding and reduced polymerase activity, ultimately hindering viral replication. These results reveal a previously unrecognized role for G4 structures in the HCV replication cycle, highlighting their importance in regulating (+) strand RNA synthesis.

**Importance:** Hepatitis C virus is a positive-sense single-stranded RNA virus that relies on an intermediate negative strand to generate new genomic RNA. While the last 157 nucleotides of the negative-strand 3’ untranslated region are known to be essential for replication, the underlying regulatory mechanisms have remained poorly understood. Our study demonstrates that a highly conserved guanine-rich sequence (107-131 nt) located on the negative strand SLIIy’ forms a guanine-quadruplex secondary structure that plays a pivotal role in orchestrating viral RNA synthesis. This G4 not only promotes efficient replication but likely acts as a molecular target to recruit the NS3 helicase, thereby allowing access for the NS5B polymerase. Disruption of the G4 structure and stabilization of the stem-loop severely impairs viral replication. These findings reveal an unrecognized layer of post-transcriptional regulation in the HCV lifecycle and establish G4 RNA structures as a critical element in HCV genome replication.

## Introduction

Hepatitis C virus (HCV) is a member of the Flaviviridae family and belongs to the Hepacivirus genus. HCV is an enveloped virus with a single-stranded positive-sense RNA genome approximately 9.6 kb in length. There are 7 HCV genotypes (>30% sequence variability) and numerous subtypes that differ in their geographic distribution, disease progression, and response to antiviral treatments(1). Hepatitis C virus (HCV) genotypes 1, 2, and 3 are the most prevalent worldwide. However, only a few clones can be grown in cell culture, such as genotype 2a (JFH-1), genotype 1a (H77), and genotype 1b (Con1(2–4). The HCV genome consists of a single open reading frame (ORF) flanked by highly structured untranslated regions (UTRs) at .he 5′ and 3′ ends, These play critical roles in translation, replication, encapsidation, and genome stability(5). The ORF encodes a polyprotein that undergoes co- and post-translational processing by host and viral proteases. This results in the formation of structural proteins (core, E1, E2) and non-structural (NS) proteins (NS2, NS3, NS4A, NS4B, NS5A, and NS5B), all of which are essential for viral replication and assembly.

RNA secondary structures are involved in almost every facet of the HCV life cycle, facilitating key processes such as translation, replication, and genome packaging (6). The 5′ UTR on the positive-sense genomic strand contains a highly conserved internal ribosome entry site (IRES) and microRNA-122 (miR-122) binding sites, which enable cap-independent translation of the viral polyprotein by directly recruiting ribosomes (7). Additional cis-acting RNA elements (CREs) are present within the UTRs and protein-coding region, contributing to RNA translation and/or genome replication (8). The 3′ UTR contains conserved stem-loop structures and poly(U/UC) sequences essential for initiating (-) strand RNA synthesis by interacting with viral and host proteins (8).

Emerging evidence suggests that guanine quadruplexes (G4s) may also be present within the HCV genome, adding another layer of regulatory complexity (9–12). G4s are highly structured nucleic acid secondary structures in guanine-rich DNA and RNA regions. These structures arise when four guanine (G) bases associate with each other through Hoogsteen hydrogen bonding, thus creating a planar G-tetrad. Multiple G-tetrads stack on top of each other, stabilized by monovalent cations such as potassium (K⁺) or sodium (Na⁺), which help maintain structural integrity. G4s can form in various topologies—parallel, antiparallel, or hybrid— depending on the strand orientation and loop connectivity (13). G4s are often predicted using software algorithms that look for a stretch of guanines 2-4 nucleotides (nt) in length, separated by a loop of 1-7 nt: G_2+_N_1+_G_2+_ N_1+_G_2+_ N_1+_G_2+_ (14).

Given that G4 structures have been shown to influence replication and gene expression in other *flaviviruses*, their potential role in HCV genome regulation warrants further investigation (15, 16). Previous studies have identified a G4 within the core (C) gene of the HCV (+) strand and another in the (-) strand 3′ UTR (9–12). Binding of the HCV core transcript G4 by cellular nucleolin or by G4-binding small molecules suppresses HCV replication and inhibits C protein translation (10, 12). Studies on the G4 located in the (-) strand 3′ UTR of HCV reveal that the G4-stabilizing molecule Phen-DC3 binds to this structure, and RNA-dependent RNA polymerase (RdRp) assays indicate that its stabilization leads to transcription elongation termination (11). Further *in vitro* studies suggest that the viral helicase NS3 can unwind these G4 structures and that their resolution is required for efficient transcription initiation (9).

During HCV replication, the RNA-dependent RNA polymerase NS5B synthesizes the positive-sense genomic RNA from the negative-strand intermediate, a process thought to be tightly regulated by RNA structural elements (17). Structural predictions and biochemical experiments suggest that within the negative-strand 3′ UTR, a specific region known as SLIIy’ forms a stable hairpin structure, which may contribute to genome stability and replication by facilitating interactions with viral and host proteins (18, 19). However, we hypothesize that the presence of a G-rich sequence within SLIIy’ may lead to the formation of a more stable guanine quadruplex (G4) rather than the hairpin described by Smith et al. (18) and may be critical in recruiting viral factors required to facilitate (+) strand RNA synthesis.

In this study, we build upon the *in vitro* observations made by Jaubert et al. (2018) and Belachew et al. (2022), both of which provided detailed evaluations of the SLIIy’ G4 and its potential role as a regulatory element (9, 11). Previous work by Friebe and Bartenschlager (2009) identified that this G4 sequence is located between nucleotides 110 and 131 and falls within the minimal region (the last 157 nucleotides of the 3’UTR) required for efficient HCV replication (19). To further investigate its functional significance, we employed a combination of bioinformatics, biophysical, and cell biology techniques. Our findings show that stabilizing the predicted hairpin structure within SLIIy’ disrupts HCV genomic RNA synthesis, while guanine-to-adenine mutations within nucleotides 110 to 131 prevent G4 formation and similarly impair RNA synthesis. We hypothesize that the G4 in the negative-strand 3′ UTR of HCV is evolutionarily conserved due to the recruitment of NS3 to unwind the viral genome and then NS5B to initiate efficient positive-strand RNA synthesis.

## Materials and Methods

### Bioinformatic analysis of the (-) strand 3’ UTR of HCV

HCV (+) strand 5’UTR sequences were obtained from the HCV Database curated by Los Alamos National Laboratory (https://hcv.lanl.gov/content/index). Under HCV sequence alignments, web alignment sequences from the 5’UTR from 2008 were accessed. The web alignment sequences have been manually optimized by the database and contain only one sequence per patient. In total, 339 sequences were obtained from genotypes 1-7: gt1 = 168, gt2 = 47, gt3 = 51, gt4 = 7, gt5 = 3, gt6 = 62, gt7 = 1. Analysis was performed in Geneious Prime to obtain consensus sequences for each genotype. The (+) strand sequences were complemented from nucleotides 110-131 to obtain the potential G4 forming sequence located on the SLIIy’ on the negative strand of HCV.

### Computational design of wild-type and mutant HCV G4 forming sequences

The consensus sequence of nucleotides 110-131 from the HCV (-) strand was evaluated using the G4Hunter algorithm to predict putative G4 structures within this region(20). The mean of the scored nucleic acid sequence was computed for a sliding window set at 20, with a threshold of 1. To investigate the role of this G4 during the HCV life cycle, mutants containing a successive introduction of guanine to adenine were generated and assessed using the G4Hunter algorithm to design constructs that lose the predicted ability to form G4s. In turn, the wild-type and mutant constructs were also evaluated using structural predictions from the RNAfold Webserver (21) with the default parameters, including G-Quadruplex formation in the structure prediction algorithm. A combination of these tools was used to generate mutants for testing using CD Spectroscopy and for functional cell culture assays.

### Oligo Purification and Folding of Oligos into G4s

Oligonucleotides at a scale of 1 µmole were ordered from Alpha DNA (Montreal, QC, Canada) and reconstituted in 1.2mL buffer containing 20 mM HEPES (pH 7.0) and 1 mM EDTA. Size-exclusion chromatography (SEC) was conducted on the Akta Pure™ system (GE Life Sciences) equipped with a Superdex 200 10/300 GL (GE Life Sciences). The column was equilibrated in 20 mM HEPES (pH 7.4), and 1 mM EDTA and 1.1 mL of the oligo was injected onto the column and isocratic elution at a rate of 0.5 mL/min and fractionation volume of 0.5 mL. The peak was collected for each oligo and utilized for downstream assays.

Following oligo purification via size-exclusion chromatography, each sample was rebuffered in 1x G4 buffer (20 mM HEPES, pH 7.4, 100 mM KCl, 1 mM EDTA) and then heated at 95°C for 5 minutes to denature any secondary and allowed to cool to room temperature over 30 minutes so that the oligos can adopt the most energetically favourable configuration. For Circular Dichroism (CD) Spectroscopy, samples were diluted to 20 µM, and for SAXS analysis, to 3 mg/mL.

### Circular Dichroism Spectroscopy

As described previously (22), Jasco J-815 spectropolarimeter (Jasco Inc.) was used to collect spectra of all oligos ranging from 220 to 320 nm, using a 1.0 mm cell, 0.1 nm data pitch, 200 µL of 20 µM folded G4 oligos sample with four accumulations and 32 s integration time. A continuous supply of nitrogen gas was provided to flush out oxygen in order to prevent ozone damage to the optics. All measurements were baseline corrected with 1x G4 buffer and repeated in triplicate.

### Small-angle X-ray scattering (SAXS)

SAXS data for all the oligos were acquired using the B21 BioSAXS beamline at the Diamond Light Source (Didcot, UK) and the Agilent 1200 (Agilent Technologies, Stockport, UK) HPLC connected to a specialized flow cell at and flowed through a pre-equilibrated SEC Shodex KW403-4F column (Showa Denko America, Inc), following previously established protocols (22, 23).

In order to ensure the homogeneity of the sample, SEC-SAXS was performed. Here, the SAXS data is collected from the sample that comes out of the HPLC. Briefly, 50 µL of 3mg/mL sample was injected into a Shodex KW402.5-4F column, pre-equilibrated with 1× G4 buffer (20 mM HEPES, pH 7.4, 100 mM KCl, 1 mM EDTA). The sample eluted out of this column was exposed to X-rays, the data was collected in frames of 3 seconds each, and a total of 600 frames were collected. Baseline correction through buffer subtraction was performed from the sample peak intensity regions using CHROMXIS (24). The buffer-subtracted scattering data were further analyzed using the ATSAS suite (version 3.0.3) (25). The Guinier approximation analysis (q² vs. ln(I(q)) was performed to evaluate sample quality and determine the radius of gyration (R_g_) (26). Structural characterization of flexibility and compactness of the wild-type and mutant oligos was assessed using a dimensionless Kratky plot (qR_g_ vs. qR_g_² * I(q)/I(0)) (27). Further, GNOM was used to generate the electron pair-distance distribution function plot; the resulting P(r) was used to determine R_g_ and maximum particle dimension (D_max_) (28). This P(r) plot was implemented in DAMMIN (29), and 10 models per oligo were independently generated (30). Finally, these 10 models were averaged using DAMAVER, and then DAMFILT was utilized to produce a filtered representative structure model (31). The models were rendered using PyMOL (32).

### Generation of replicon mutants

The pFK341-Sp-PI-luc-EI-core-3′JFH1 replicon system (Gift from Dr. Ralf Bartenschlager) developed by Friebe and Bartenschlager (2009) was used for all replicon assays (19). This reporter virus system contains the HCV 5’UTR followed by a 61-nt-long spacer element, then a poliovirus IRES, firefly luciferase (Fluc) reporter gene, Encephalomyocarditis virus IRES and then the HCV JFH-1 coding sequence followed by the 3’UTR. 1µg G-blocks containing the G4 mutants **(see Table 1 for sequences)** ordered from Twist Biosciences were resuspended in 100 µL of water. PCR was performed according to manufacturer protocols using Phusion polymerase (M0530, NEB, Ipswich, MA, USA) and was conducted using the following primers purchased from IDT: HCV_Sbf1_FWD: TAAGCTGCCTGCAGGTAATACGAC and HCV_PmeI_REV: ACGGGGGTTTAAACGGTGCA. The PCR amplification protocol was as follows: initial denaturation at 98° C for 30 seconds, 35 cycles of denaturation at 98° C for 10 seconds, annealing at 67.4° C for 30 seconds, and extension at 72° C for 15 seconds, concluding with a final extension at 72° C for 10 minutes. PCR cleanup was performed using the EZNA cycle pure kit (D6492, Omega Biotek, Norcross, GA, USA), and the amplified DNA products were linearized overnight using SbfI-HF and PmeI enzymes (R3642L and R0560L, NEB, Ipswich, MA, USA). After purification using the EZNA Cycle Pure Kit, the digested products were ligated into the SbfI/PmeI-digested pFK341-Sp-PI-luc-EI-core-3′JFH1 wild-type and replication-dead ΔGDD backbones. Prior to ligation, these backbones were treated with Quick Calf Intestinal Alkaline Phosphatase (Quick CIP, M0525L, NEB, Ipswich, MA, USA) and PCR cleanup was performed. The ligation, using T4 DNA Ligase (M0202L, NEB, Ipswich, MA, USA), was performed overnight at 16° C. The ligated products were transformed into *E. coli* NEB5α cells (C2987H, NEB, Ipswich, MA, USA) and positive colonies were selected on LB-ampicillin agar plates. Positive colonies were further grown in LB + ampicillin media, and DNA was isolated using the EZNA plasmid mini kit II (D6945-00, Omega Biotek, Norcross, GA, USA). Correct gene insertion into the plasmid DNA was confirmed via Sanger sequencing at Eurofins Genomics (USA) using the HCV_Sbf1_FWD primer.

**Table 1.**
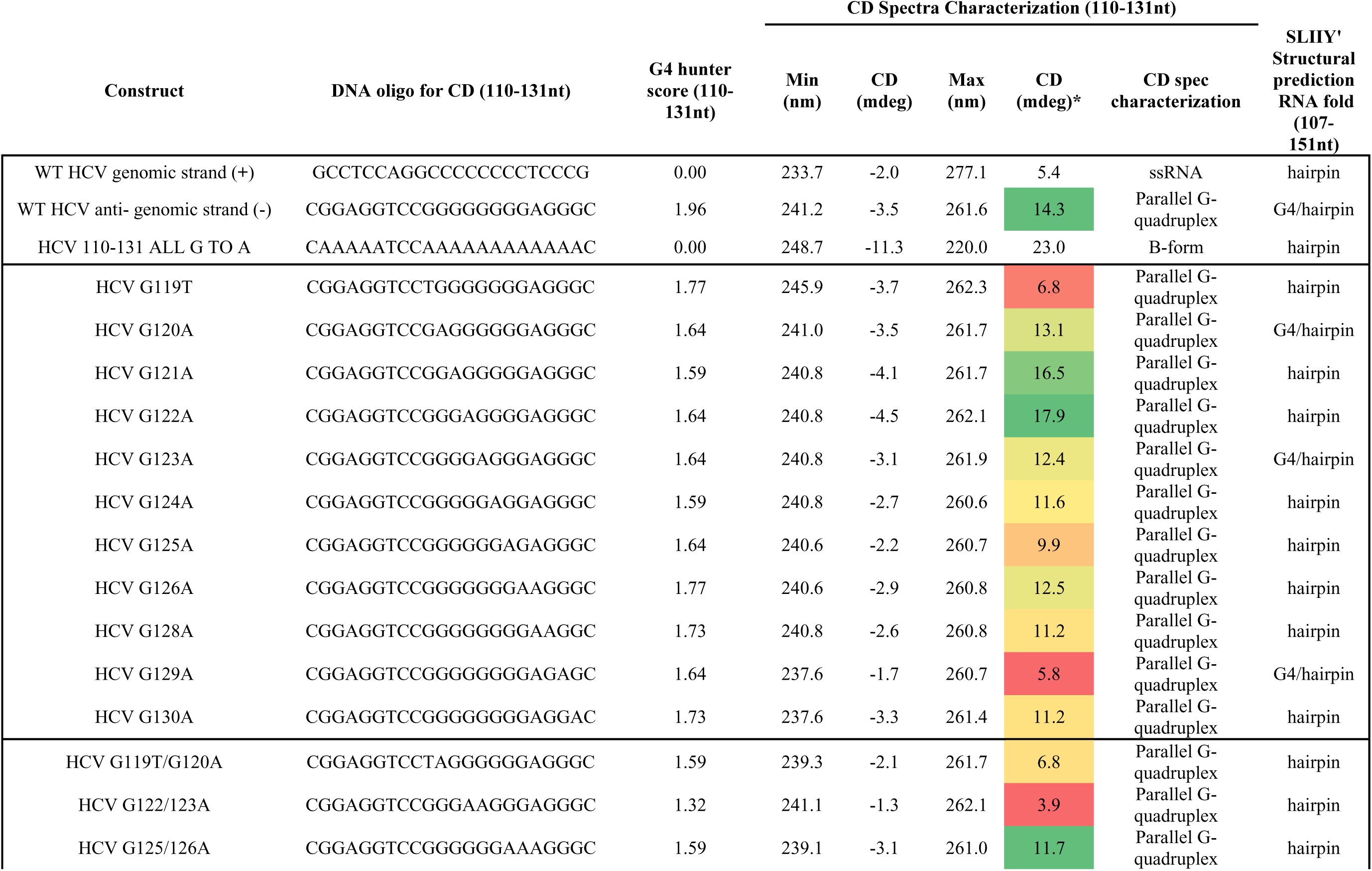
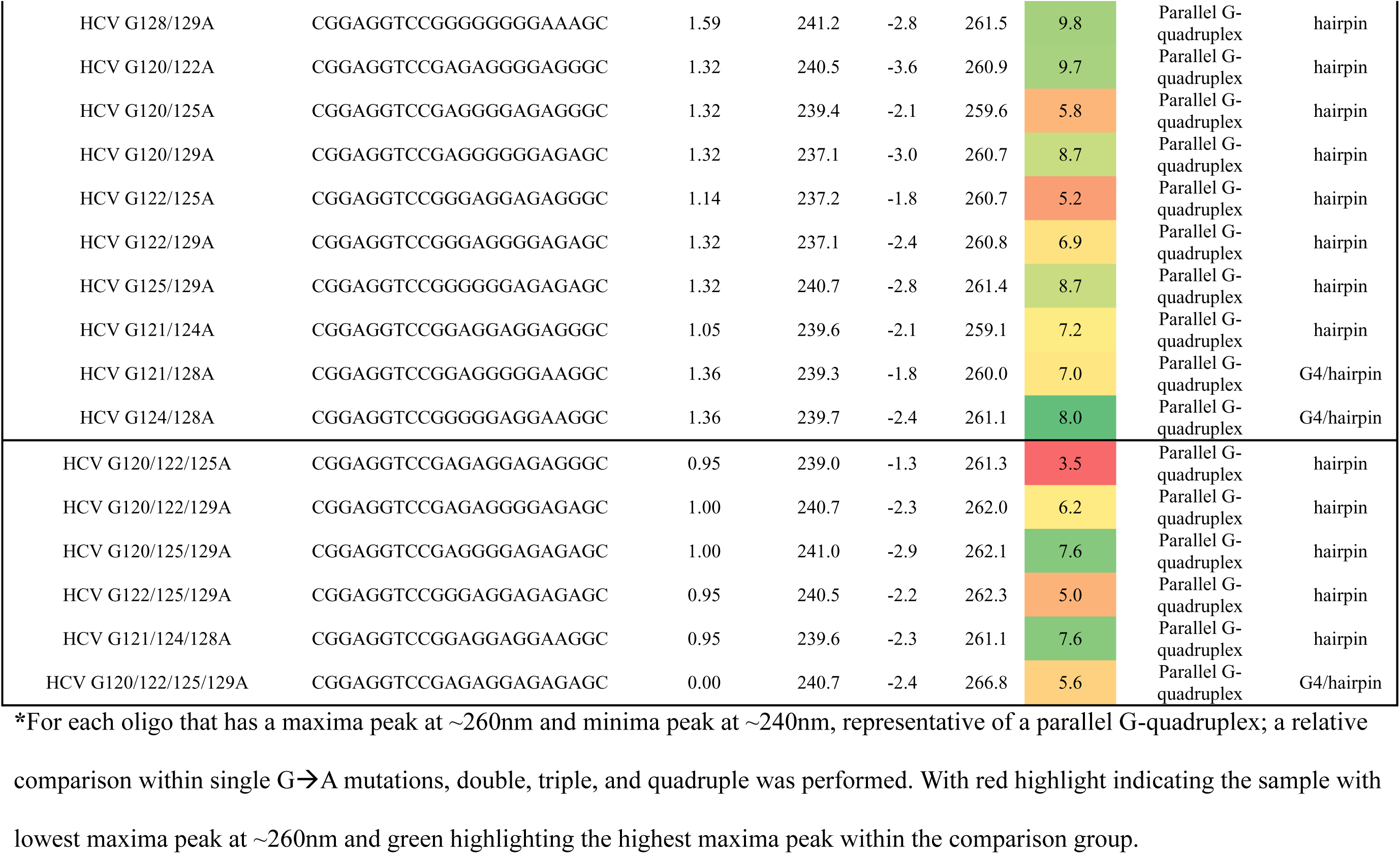
Construct design for HCV oligos characterized using G4 hunter, RNAfold, and CD-spectroscopy.

### HCV RNA *in-vitro* transcription

For IVT preparation, the sequence confirmed wild-type, mutant, and replication-dead pFK341-Sp-PI-luc-EI-core-3′JFH1 plasmid constructs were grown in NEB5α cells but prepped using the Endo-free plasmid DNA mini kit II (D6950-00, Omega Biotek, Norcross, GA, USA). Plasmids were linearized using the MluI-HF (R3198L, NEB, Ipswich, MA, USA) restriction enzyme overnight at 37° C. A phenol-chloroform-isoamyl alcohol extraction was performed, and the DNA was precipitated with 3 M 0.1 CV sodium acetate and 2.5 CV of 100% ethanol. The sample was stored at -80° C overnight, washed with 70% ethanol, and resuspended to achieve a final concentration of 1 µg/µL.

In a 100 µL reaction volume, 2 µg of plasmid DNA was added along with 1 mM ATP/CTP/UTP and 1.2 mM GTP (R0481, Thermo Fisher Scientific, Waltham, MA, USA), 0.8 U/µL RiboLock (EO0382, Thermo Fisher Scientific, Waltham, MA, USA), and 200 U T7 RNA polymerase (M0251L, NEB, Ipswich, MA, USA). These components were combined and incubated at 37° C for 3 hrs. DNaseI (RQ1, Promega, Madison, WI, USA) was then added at 2 U/µg plasmid DNA and incubated at 37° C for 20 minutes. On a formaldehyde-denaturing agarose gel, 2 µL of the IVT was run to ensure that the RNA was intact. To stop the DNaseI digestion, the IVT RNA was precipitated using 1 µL GlycoBlue™ co-precipitant (AM915, Thermo Fisher Scientific, Waltham, MA, USA), 0.1 CV 3 M sodium acetate, and 2.5 CV of 100% ethanol. The RNA was stored at -80° C until ready to use. On the day of use, the RNA was precipitated, washed with 70% ethanol, and then resuspended in 100 µL of nuclease-free water and nano-dropped.

Capped renilla luciferase control, RNA was prepared using the pRL-SV40 Vector (E2231, Promega, Madison, WI, USA). The vector was linearized overnight at 37° C using the HpaI (R0105L, NEB, Ipswich, MA, USA) restriction enzyme and precipitated as previously described. RNA was prepared in a 100µL reaction volume using the mMESSAGE mMACHINE T7 Transcription Kit (AM1344, Invitrogen, Waltham, MA, USA) and treated with DNAse, then precipitated as outlined above for the HCV RNAs.

### Cell Culture

The protocol used was obtained from Rheault et al. (7)with slight modifications described below. For simultaneous luciferase assays with RT-qPCR, two 0.4 cm cuvettes (Bio-Rad Laboratories, Hercules, CA, USA), each containing 8 x 10^^6^ Huh7.5 cells (gift from Prof. Stephan Urban), were resuspended in 400 µL of cold phosphate-buffered saline (14190144, Thermo Fisher Scientific, Waltham, MA, USA) and mixed with 20 µg of HCV-Fluc RNA and 2 µg of RLuc RNA. These were electroporated at 270 V, 950 μF, and infinite resistance in the Bio-Rad Gene Pulser XCell (Bio-Rad Laboratories, Hercules, CA, USA). The two cuvettes were then combined into 9.2 mL of media. For the luciferase assays, cells were plated in 12-well plates with 300 µL for the 4-hour time point, and 150 µL for the 24–72-hour time points. For harvesting RNA, the cells were plated in 6-well plates with 1000 µL for the 4h time point and 500 µL for the 24–72-hour time points. Media was topped up in the 12-well and 6-well plates to reach a final volume of 1 mL and 2 mL, respectively.

### Luciferase Assays

For the luciferase assays, cells were washed in PBS and harvested in 200 μL of freshly diluted 1× passive lysis buffer (E1941, Promega, Madison, WI, USA). Cells were incubated on a shaking platform for 15 minutes, scraped with the back of a pipette tip, collected, and frozen at - 80° C until all time points were collected. Luciferase assays were performed using the Dual-Glo Luciferase Assay System (E2920, Promega, Madison, WI, USA). Briefly, the samples and assay buffers were equilibrated to room temperature, and 25 μL of sample was aliquoted in a white 96-well plate (CLS353296, Corning, Corning, NY, USA) and mixed with 25 μL of luciferase assay reagent, incubated for 10 minutes, and read on a Promega Glomax Multi Detection System luminometer (Promega, Madison, WI, USA) with an integration time of 10 seconds, read in duplicate. Following the luciferase readout, Renilla was measured by adding the Stop & Glo substrate, incubating for 10 minutes, and reading.

Renilla luciferase was used only as a transfection control and not for data processing. Data was normalized to the input of the highest RLUs of firefly luciferase detected at the 4-hour electroporation time point to account for any differences in electroporation variability.

### HCV RNA quantification

Cells in 6-well plates were first washed with PBS and incubated for 15 minutes on a shaking platform with 250 μL TRIzol reagent (Thermo Fisher Scientific, Waltham, MA, USA), then transferred to -80° C until samples were ready for extraction. The total RNA was extracted according to the manufacturer’s instructions, resuspended in 20 µL of nuclease-free water, and subjected to nanodrop analysis. One-step RT-qPCR for quantification of total HCV RNA, as described by Rheault et al., was performed on the Biorad C1000 Touch CFX96 Real Time (Bio-Rad Laboratories, Hercules, CA, USA) system with slight modifications(7). The total reaction volume for RT-qPCR was 10 μL, containing 0.4 μM of NS5B-FOR (5′-AGA CAC TCC CCT ATC AAT TCA TGG C-3′) and NS5B-REV (5′-GCG TCA AGC CCG TGT AAC C-3′) primers, 0.2 μM of the NS5B-FAM-BHQ1 probe (5′-ATG GGT TCG CAT GGT CCT AAT GAC ACA C-3′) (AlphaDNA, Montreal, QC, Canada), 200 ng of RNA, 25 U Maxima H Minus reverse transcriptase (EP0753, Thermo Fisher Scientific, Waltham, MA, USA), and the iTaq Universal Probes Supermix (Bio-Rad Laboratories, Hercules, CA, USA). The protocol for RT-qPCR is as follows: 55° C for 30 mins, 95° C for 2 mins, and 45 cycles of 95° C for 15 seconds and 64° C for 30 seconds. Genome copies were calculated against a genomic RNA standard curve. Data was normalized to the highest amount of RNA detected at the 4-hour electroporation time point to account for any differences in electroporation variability.

## Results

### Bioinformatic analysis of the negative strand 3’UTR

The last 157 nucleotides at the 3’ end of the HCV (-) strand play a critical role in initiating (+) strand genomic RNA synthesis. Jaubert et al. (2018) previously identified a guanine quadruplex (G4) between nucleotides 110–131 using the G4Hunter algorithm. To build on these findings, HCV 5’UTR sequences were retrieved from the HCV Database curated by the Los Alamos National Laboratory (https://hcv.lanl.gov/content/index). Under the HCV Sequence Alignments section, web-aligned sequences from the 5’UTR dating back to 2008 were accessed. These alignments have been manually optimized to include only one sequence per patient. In total, 339 sequences from genotypes 1–7 were obtained **(see Supplementary S1):** gt1 = 168, gt2 = 47, gt3 = 51, gt4 = 7, gt5 = 3, gt6 = 62, gt7 = 1. A web logo alignment was generated from these sequences **(Fig 1A)**, revealing variability at nucleotide position 119, where uracil is most commonly found, followed by guanine. The remaining positions appear to be highly conserved across all genotypes.

**Figure 1.**
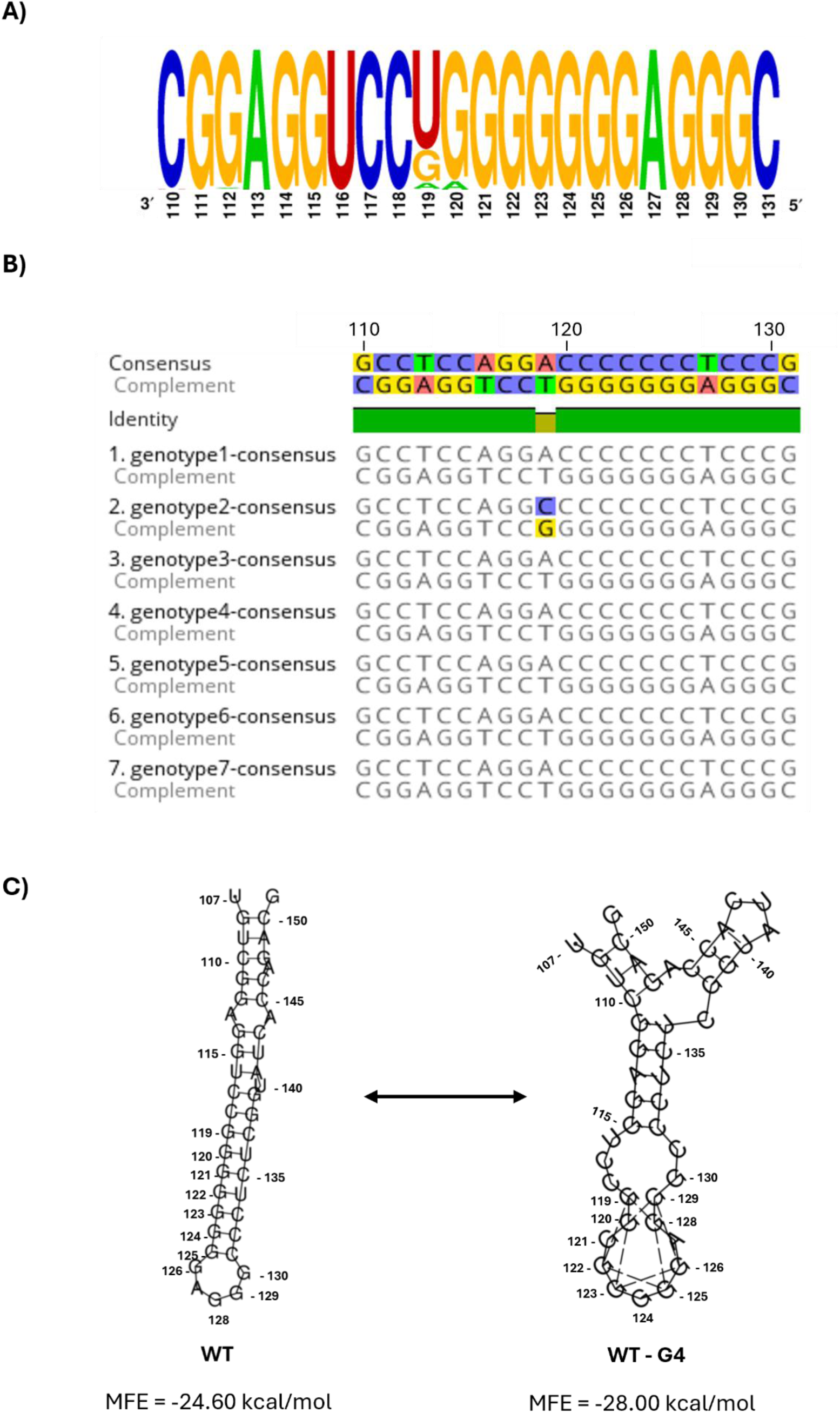
Computational assessment of nucleotides HCV nucleotides 110-131. **(A)** A total of 338 sequences of the (+) strand HCV 5’UTR nucleotides 110-131 from genotypes 1-7 were retrieved from https://hcv.lanl.gov/. Sequences were then complemented to obtain the sequences of the (-) strand and aligned using WebLogo application. (**B)** Consensus sequence of nucleotides 110-131 in the 5’ UTR of the (+) strand and the 3’ UTR of the (-) strand of HCV, obtained by aligning individual genotypes 1-7 using Geneious Prime. Guanine is frequently found at position 119 in genotype 2, whereas thymine/uracil is in all other genotypes. (**C)** RNAfold computationally predicted secondary structure of the SLIly’ (nucleotides 107-151) in the (-) strand 3’ UTR of genotype 2a JFH1, generated with and without the G4 prediction algorithm.

To further investigate genotype-specific preferences at position 119, the cDNA clones obtained from the HCV Database were analyzed using Geneious Prime to generate consensus sequences for each genotype **(Fig 1B)**. The (+) strand sequences were then complemented from nucleotides 110–131 to identify the potential G4-forming sequence located on the SLIIy’ region of the negative strand. Interestingly, genotype 2 exhibited a preference for guanine at position 119, whereas all other genotypes predominantly contained uracil. This observation is notable, as JFH-1, the strain commonly used in cell culture and replicon assays, is derived from genotype 2a. Consequently, guanine was selected at position 119 for structural analysis using the RNAfold Webserver, run with default parameters and structures were rendered with the option to incorporate G4 in the structural prediction algorithm selected (**Fig 1C, right panel)** or deselected **(Fig 1C, left panel)**. While the G4Hunter algorithm identifies G4 motifs solely based on sequence composition, RNAfold was included to assess whether the software could also predict G4 formation based on sequence and structural features. In addition to structure prediction, RNAfold calculates the Minimum Free Energy (MFE), the lowest possible Gibbs free energy (ΔG), representing the energy of the most thermodynamically stable structure a given nucleic acid sequence can adopt. The MFE for the canonical hairpin structure, which is widely accepted within the field, was found to be -24.6 kcal/mol, whereas the predicted G4 structure had an MFE of -28 kcal/mol. Since a more negative MFE indicates greater thermodynamic stability (33), these results suggest that the G4 conformation is more energetically favourable than the traditional hairpin structure.

To computationally assess the stability of the putative G4 structures within this region, the consensus sequence for genotype 2a nucleotides 110–131 from the HCV (-) strand was analyzed using the G4Hunter algorithm. The mean of the scored nucleic acid sequence was computed for a sliding window arbitrarily set at 20, with a threshold of 1. A potential G-Quadruplex-Forming Sequence (PQS) is given for each sequence, where a score >1 indicates possible G4 formation and a score of <1 indicates unlikely to form a G4. To evaluate the function of this G4 during the HCV life cycle, HCV (-) strand mutants within the PQS at nucleotides 110– 131 were generated by successively replacing guanosines with adenines. These mutants were designed using the G4Hunter algorithm to disrupt the predicted ability to form G4 structures **(Table 1)**. Additionally, the wild-type and mutant constructs were evaluated using structural predictions of the RNAfold Webserver with the default parameters and the incorporated G-Quadruplex formation into the structure prediction algorithm. A combination of these tools was used to generate mutants to test using CD Spectroscopy and for the functional cell culture assays.

### Biophysical characterization of WT and Mutant HCV oligos – CD Spectropolarometry

As shown in **Table 1**, 33 constructs spanning nucleotides 110–131 were designed to assess their ability to form a guanine quadruplex using biophysical methods. The panel included 10 single guanine-to-adenine (G→A) mutants, 1 guanine-to-thymine (G→T) mutant to represent the 119T sequence found in all genotypes except for genotype 2, which is 119G; 13 double mutants; 5 triple mutants; and 1 quadruple mutant. Additionally, the wild-type (WT) 110–131 (+) strand sequence was included as a negative G4 control, along with a mutant in which all guanines were replaced with adenines (HCV 110–131 all G→A). DNA oligonucleotides were used for G4 characterization instead of RNA due to their stability and provided comparable data for HCV 110-131 to the RNA oligos utilized by Jaubert et al(11). Each DNA oligo was purified via size-exclusion chromatography in a 20 mM HEPES (pH 7.4) and 1 mM EDTA buffer to ensure that a homogenous preparation of the oligo could be obtained. All chromatography plots for the oligo purification can be found in **Supplementary Figure S3**. All oligos had a peak between 13-14 mL elution volume, and these samples were collected for downstream analysis.

Guanine quadruplexes require the presence of a monovalent cation to form. G4s are best stabilized by potassium ion (K⁺), which can coordinate with the electronegative carbonyl groups in the G-tetrad planes, promoting the formation and structural integrity of these secondary structures (34). Notably, intracellular conditions are particularly favorable for G4 formation, as the cytoplasm typically contains 100-150 mM of potassium ions, creating an optimal ionic environment for G4 stability. Heat-cooling, or thermal annealing, is commonly used for *in vitro* preparation of G4 oligonucleotides because it helps promote the proper folding of these structures into their most thermodynamically stable conformations. As such, the peak from size exclusion chromatography for each oligo was collected (∼ 0.5 mL), and the volume was adjusted to prepare a final concentration of 20 µM in 1x G4 buffer (20 mM HEPES pH 7.4, 100 mM KCl, 1 mM EDTA). All samples were heated at 95°C for 5 minutes and then slowly cooled to room temperature over the course of 30 minutes.

The ellipticity was measured for each sample in the G4 buffer from 220 nm to 320 nm. **Figure 2A** shows the CD spectroscopy profiles of the WT HCV (+) and (-) strands, along with the complete G→A mutant. The WT HCV (+) strand (nucleotides 110–131) exhibits a broad positive peak at 280 nm with a negative peak at 233 nm, consistent with single-stranded DNA (ssDNA) formation. The mutant HCV 110–131 all G→A displays a negative peak at 248.7 nm and a positive tail between 220 and 240 nm, characteristic of B-form DNA (35). In contrast, the WT HCV (-) strand (110–131 nt) shows a negative peak of near 240 nm and a strong positive peak of 14.3 mdeg around 260 nm, which is a signature CD profile of a parallel G-quadruplex (36). Single mutants are compared to the WT oligo in **Figure 2B**. The G119T oligo, which represents the consensus sequence for all genotypes except for genotype 2, still has a spectrum profile of a parallel G4 but a slight left spectra shift with a maxima at 262.3nm and a minima at 245.9 nm. G121A and G122A had higher peak ellipticity than the WT (-) strand HCV construct, indicating that these mutations stabilized G4 confirmation further. While G129A (5.8 mdeg) and G119T (6.8 mdeg) had the lowest positive peak ellipticity, suggesting that these single-nucleotide mutations appear to strongly disrupt quadruplex formation. Based on the location of both of these single mutants in **Figure 1C**, it is possible that these nucleotides have a critical role in stabilizing the G4 structure and are found in the same position but on opposite sides of the hairpin/G4. In **Figure 2C** the double mutants all still retain the signature parallel G4 formation but have considerably lower positive peaks compared to the wild-type and single mutants, with the following mutants showing the lowest peak ellipticity: G122/123A (3.9 mdeg) < G122/125 (5.2 mdeg) < G120/125 (5.7 mdeg) < G122/129A (6.9 mdeg) = G119T/G120A (6.8 mdeg). The triple mutants and quadruple mutants in **Figure 2D** still retain their signature parallel G4 spectra, but their peaks are very low, ranging from 3.5 mdeg to 7.6 mdeg, indicating that there are a large number of quartets disrupted.

**Figure 2.**
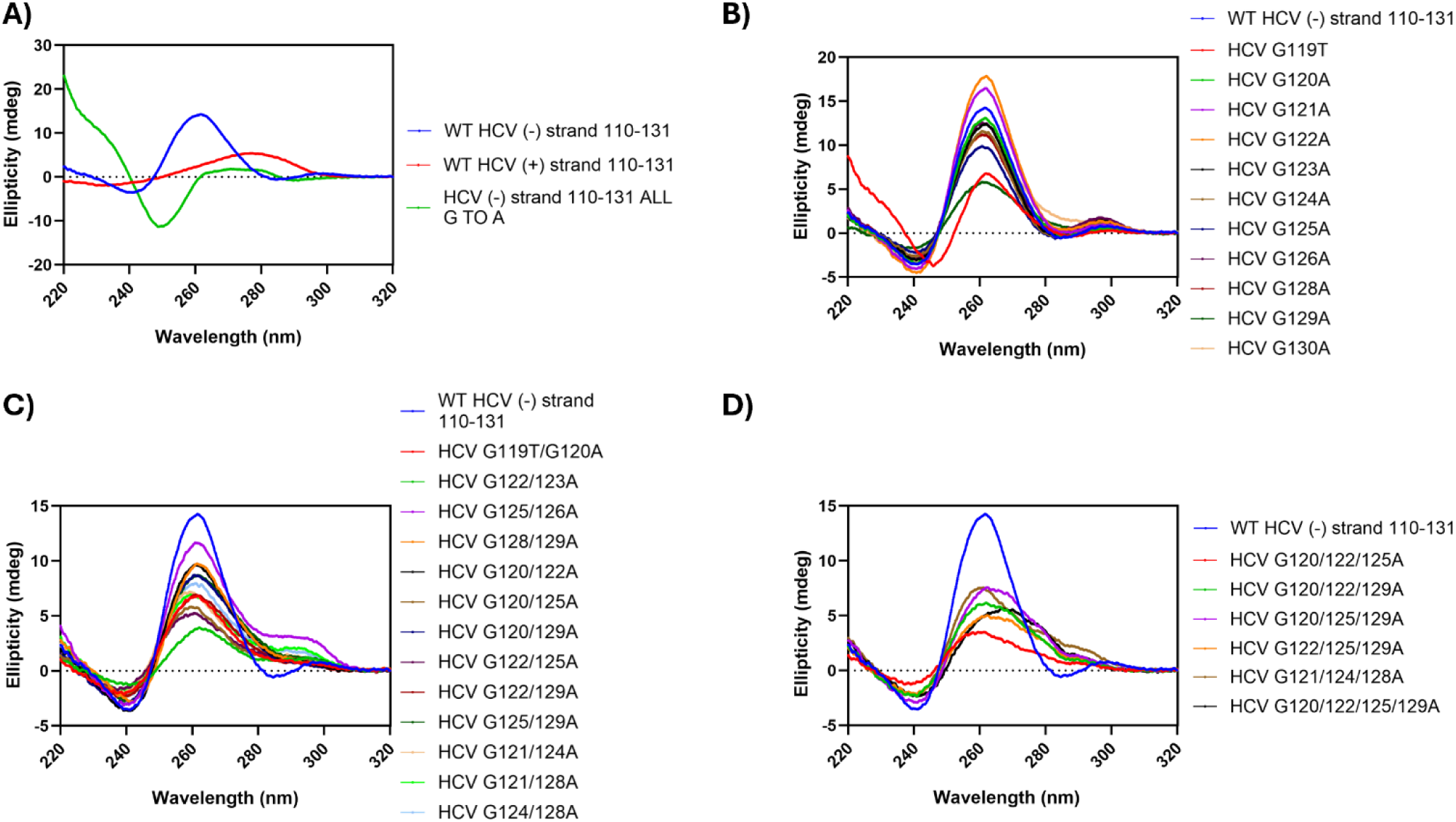
WT and mutant HCV oligos (110-131nt) are purified to homogeneity and form a G4 parallel G4 in the presence of K^+^ ions. Size-exclusion chromatography purified HCV oligos were rebuffered into G4 buffer containing 20mM HEPES pH 7.4, 100mM KCl, and 1mM EDTA, heated to 95°C for 5 minutes and slow-cooled to room temperature over 30 minutes. Circular dichroism (CD) spectra were collected for **A)** WT (+) and (-) strand HCV oligos **B)** single mutant guanine to adenine (-) strand oligos **C)** double mutant guanine to adenine, (-) strand oligos and **D)** triple and quadruplex guanine to adenine mutant (-) strand oligos. The WT (-) strand HCV oligo forms a parallel G4, as indicated by a positive peak at 265nm and a negative signal at approximately 240nm. Three biological replicates were performed for each oligo and the mean was plotted for each sample.

### Biophysical characterization of WT and Mutant HCV oligos - Small-Angle X-ray Scattering (SAXS)

We employed SAXS to visualize the low-resolution three-dimensional structures of the wild-type positive and negative strands and the negative strand mutant HCV G4 oligos (110-131nt) folded in G4 buffer. HPLC-SAXS was used to separate the molecules in a size-exclusion chromatography system prior to data collection. A single sharp peak was observed for each sample. By subtracting the scattering information from the buffer and merging the SAXS intensity plots **(Supplemental Figures S4-S34),** we could select a data set from a region with a uniform distribution. The intensity plot together with the linearity observed in the low q values of Guinier analysis **(Supplemental Figures S4-S34)** indicated that all oligos have a uniform population and are free of aggregation in G4 buffer, as also demonstrated by SEC purification **(Supplemental Figures S4-S34)**. Additionally, Guinier analysis provided valuable measurements, including the intensity (Guinier I(0)), the radius of gyration (R_g_), and the angular range (q.R_g_ range) in reciprocal space, as shown in **(supplemental figure S4-S34)**.

In several mutants, an increase in R_g._ was observed as compared to that of wildtype (WT), especially in the case of successive G→A mutations. For the WT, an R_g_ of 16.90 Å was observed, which then increased to a value of 17.30 Å for G120/122/125A and 19.52 Å for G120/125/129A, demonstrating a more elongated molecule. Following Guinier’s analysis, a P(r) analysis was performed to gain further insight into the folded state of the oligos. An indirect Fourier transformation was used to derive the P(r) plot to convert the reciprocal information on the Guinier plot into a real-space paired electron distribution. The values, as seen in **Table 2**, show only a slight variation in the reciprocal space R_g_ (Guinier R_g_ Å) and the real space R_g_ (P(r) R_g_ Å), with most cases being less than 0.1 Å. Furthermore, with the P(r) plot, we gain information about the oligo’s maximum dimension (D_max_), shown in **Table 2**, which shows how elongated that oligo is. The WT has a D_max_ of 46.35Å, and a similar trend is observed with successive G→A mutations, leading to an increased D_max_. This suggests an elongation of the molecule, exemplified by G120/122/125A and G120/125/129A, which have Dmax values of 58.48 Å and 63.14 Å, respectively. The P(r) file was implemented in DAMMIN to generate 10 independent low-resolution models of each oligo, each with a statistically favourable χ^2^value shown in **Table 2**. The 10 models were averaged using DAMAVER, and DAMFILT models were generated and presented in **Figure 3**. We also calculated the normalized spatial discrepancy (NSD) for the 10 models shown in **Table 2**, and they indicated that the DAMMIN-derived structures were in good agreement with each other.

**Figure 3.**
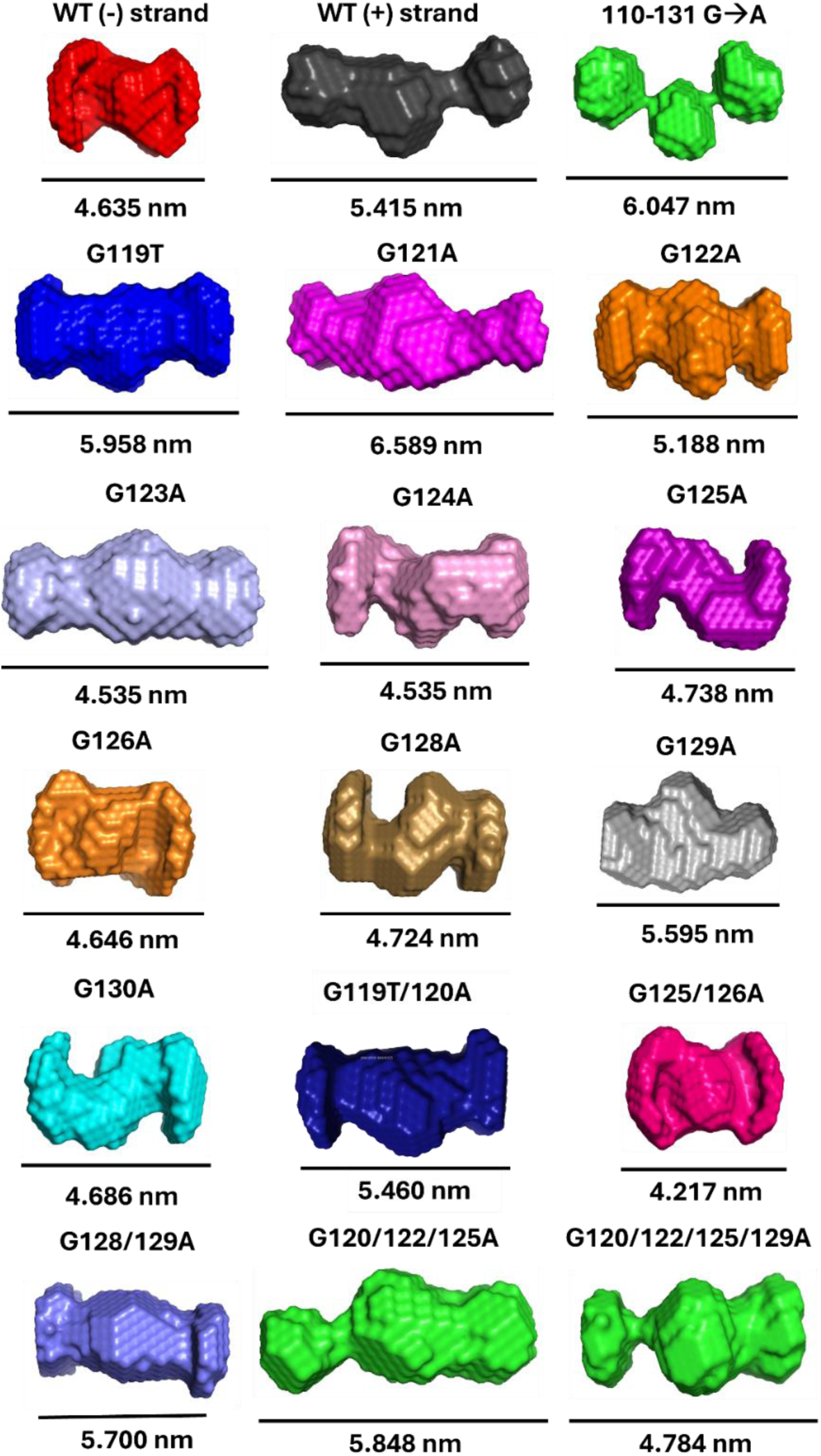
Low-resolution three-dimensional structures *ab initio* model reconstructions of WT and mutant (-) strand HCV oligos (110-131 nt) obtained from Small-Angle X-ray Scattering (SAXS). Size-exclusion purified oligos were rebuffered in 20 mM HEPES pH 7.4, 100 mM KCl, and 1 mM EDTA, heated to 95° C for 5 minutes and slow-cooled to room temperature over 30 minutes and then subjected to SAXS. The low-resolution *ab initio* models show the average electron density of each molecule in solution. The HCV WT (-) strand oligo has a compact and symmetrical structure, characteristic of a G4-like molecule with a D_max_ of 4.635 nm. Conversely, the (+) strand has an asymmetrical, elongated, dumbbell-like structure with a D_max_ of 5.415 nm. Successive guanine to adenine mutations disrupt the overall compact G4-like SAXS envelope, leading to elongated multidomain envelopes G120/122/125A D_max_ 5.848 nm and G120/122/125/129A D_max_ 4.784 nm. Complete disruption of the G4 in the (-) strand in the 110-131nt guanosine to adenine mutant appears to form a three-lobed structure connected by flexible linkers with an overall maximum dimension (D_max_) of 6.047 nm.

**Table 2.**
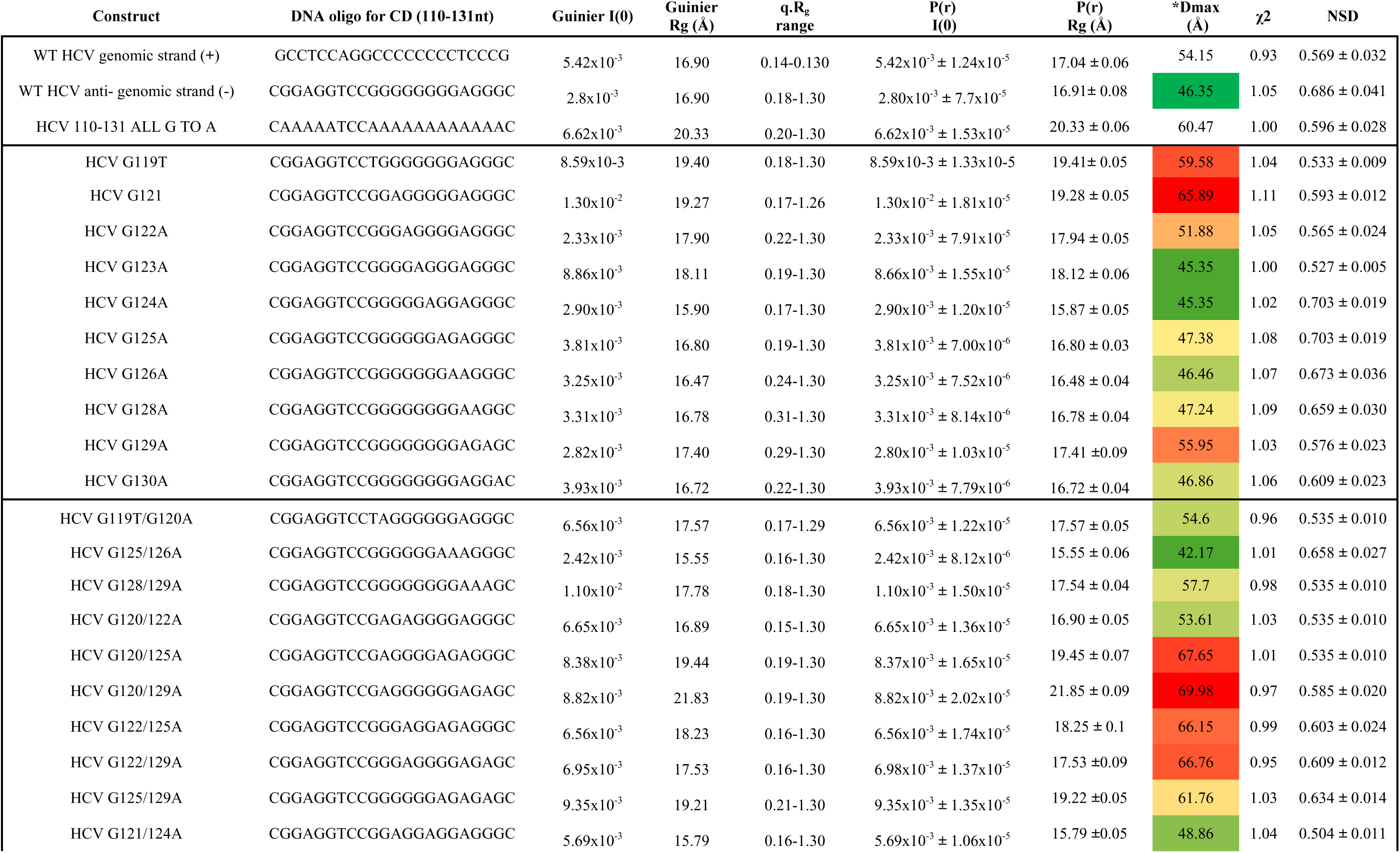
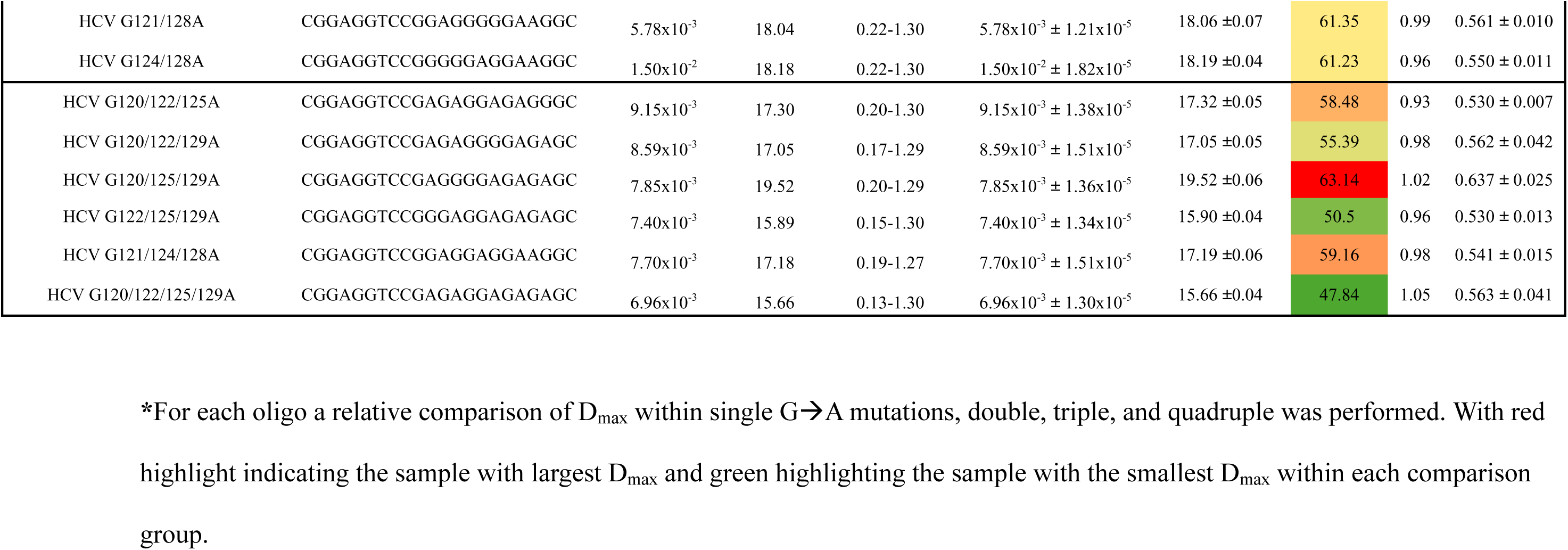
SAXS measurements and parameters of folded HCV oligos.

The representative model for each oligo used for cell culture assays is shown in **Figure 3**, with the SAXS envelopes for the additional oligos, which can be found in the **Supplementary Figures S4-S34**. The HCV WT (-) strand oligo has a compact and symmetrical structure, characteristic of a G4-like molecule with a D_max_ of 4.635nm. This wild-type G4 oligo has a similar SAXS envelope to what we have previously published for HBV and MPXV G4s (22, 23, 37). Conversely, the (+) strand appears to have an asymmetrical, elongated, dumbbell-like structure with a D_max_ of 5.415 nm. Successive guanine to adenine mutations disrupt the overall compact G4-like SAXS envelope, leading to elongated multidomain envelopes G120/122/125A D_max_ 5.848 nm and G120/122/125/129A D_max_ 4.784 nm. Complete disruption of the G4 in the (-) strand in the 110-131 nt guanosine to adenine mutant appears to form a three-lobed structure connected by flexible linkers with an overall D_max_ of 6.047 nm. These findings demonstrate that when the G4 is present, it presents as a compact structure, as when mutations disrupt it, the structure begins to elongate and, in some cases, form ball-like structures. This observation is consistent to what we have seen previously for MPXV (37).

### Investigating the role of G-quadruplex structures in HCV replication in cell culture

Traditional approaches for studying G-quadruplexes (G4s) in the HCV genome, such as chemical stabilization or inhibition using small molecules, present significant limitations. Many G4-targeting drugs, including pyridostatin and BRACO-19, exhibit broad and non-specific activity, affecting multiple G4 structures throughout the host and viral genome rather than targeting a single site. It is challenging to distinguish direct effects on the SLIIy’ G4 from secondary effects on other G4-containing regions. Another common approach is introducing mutations directly into the HCV genome, which poses additional challenges. Since G4s often overlap with functionally essential regions of the viral RNA, such as the internal ribosome entry site (IRES) or replication elements, mutations intended to disrupt G4 formation can also interfere with translation and replication. This makes it difficult to distinguish whether observed changes in viral replication result from G4 loss or structural perturbations affecting other aspects of the viral life cycle.

To overcome these challenges, we utilized an HCV replicon model developed by Friebe and Bartenschlager (2009) (19). As shown in **Figure 4**, this replicon consists of the HCV 5′ UTR, followed by a 61-nucleotide spacer, the poliovirus internal ribosome entry site (P-I), the firefly luciferase (Fluc) reporter gene, and the Encephalomyocarditis virus (EMCV) IRES (E-I), which facilitates translation of the HCV coding sequence, ending with the HCV 3′ UTR. Since the HCV polyprotein in this replicon is no longer dependent on the HCV IRES for translation, the 5′ UTR instead functions as a promoter for luciferase translation. Additionally, in the negative strand, the 3′ UTR serves as a regulatory element for assessing the impact of mutations on (+) genomic RNA synthesis. Luciferase activity is measured as a surrogate marker for (+) strand RNA accumulation, providing a quantitative readout of viral replication efficiency. The replicon system enables precise assessment of the effects of G4-disrupting mutations on viral RNA replication without the confounding influences of viral entry, assembly, or release. By selectively introducing guanine-to-adenine substitutions in the SL-IIy’ region, we have the ability to specifically evaluate the role of G4s in HCV replication while minimizing unintended disruptions to other viral processes.

**Figure 4.**
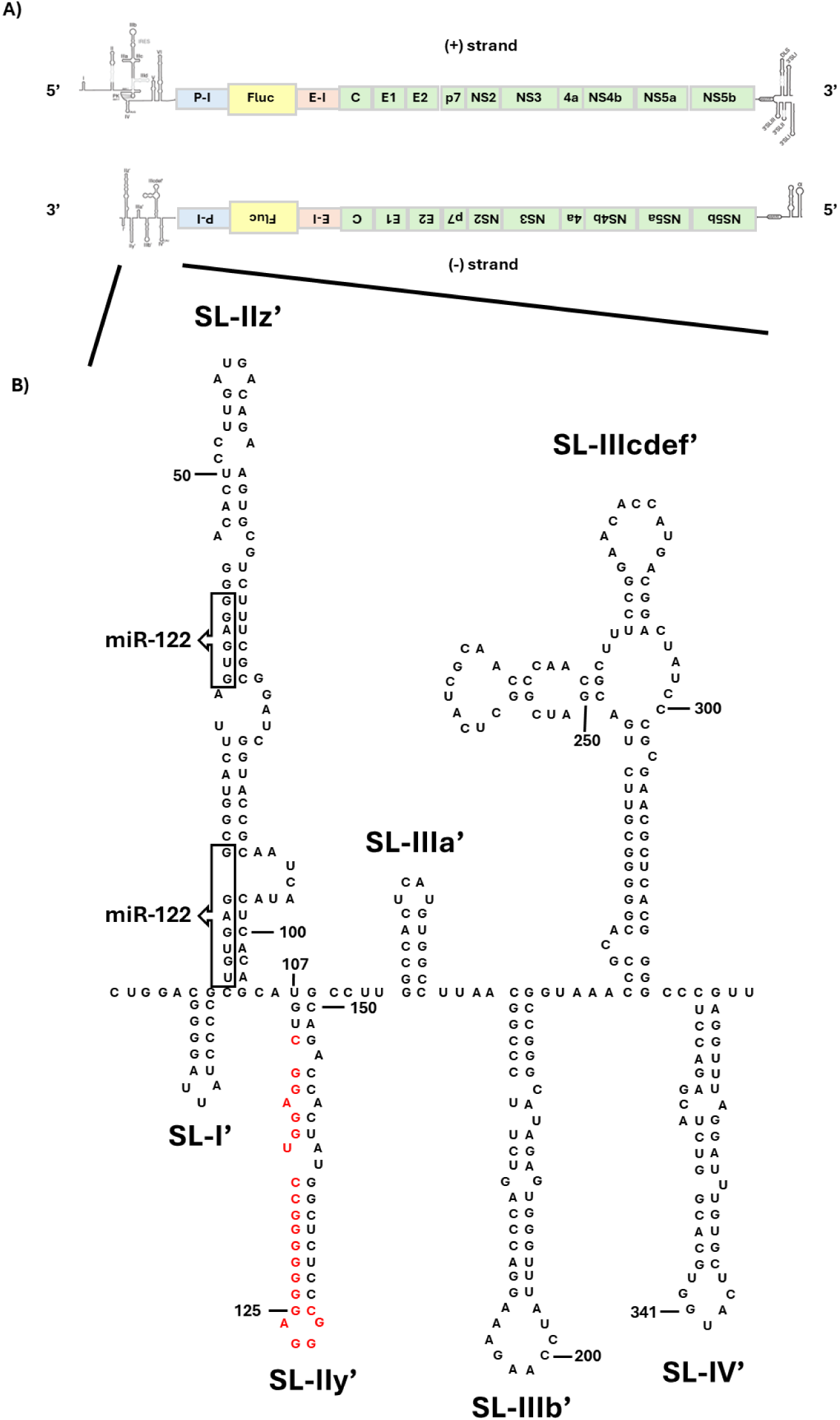
HCV replicon used to assess the role of the G4 motif in SL-IIy’ during viral replication. **(A)** Schematic representation of the JFH1-derived reporter virus genome, shown in both (+) and (−) orientations, highlights the secondary structures at the RNA’s 5′ and 3′ ends. A 61-nucleotide spacer follows the HCV 5′ UTR, the poliovirus internal ribosome entry site (P-I), the firefly luciferase (Fluc) reporter gene, and the Encephalomyocarditis virus (EMCV) IRES (E-I), which facilitates translation of the HCV coding sequence, ending with the HCV 3′ UTR. **(B)** Predicted secondary RNA structure at the 3′ end of JFH1 (−) RNA, based on Smith et al. (2002) (18), with Stem-loop (SL) structures labelled according to established nomenclature. Nucleotides are numbered from 3′ to 5′, and boxed arrows indicate complementary regions corresponding to miRNA-122 seed sequences in the 5′ NTR of (+) RNA. The G4 sequences under investigation (nucleotides 110–131) are highlighted in red. The figure was adapted from Friebe and Bartenschlager (19).

A subset of 19 out of 33 mutations previously analyzed using biophysical methods (CD spectroscopy and SAXS) was selected for testing in the replicon assays. Additionally, we included compensatory mutants (labelled COMP) that preserve the original SL-IIy hairpin structure while introducing slight sequence modifications. To evaluate the impact of these mutations on RNA secondary structures, we used RNAfold to predict structural changes and designed constructs accordingly to assess their effects on primary sequence integrity, G4 formation, and hairpin stability **(Figure 5)**.

**Figure 5.**
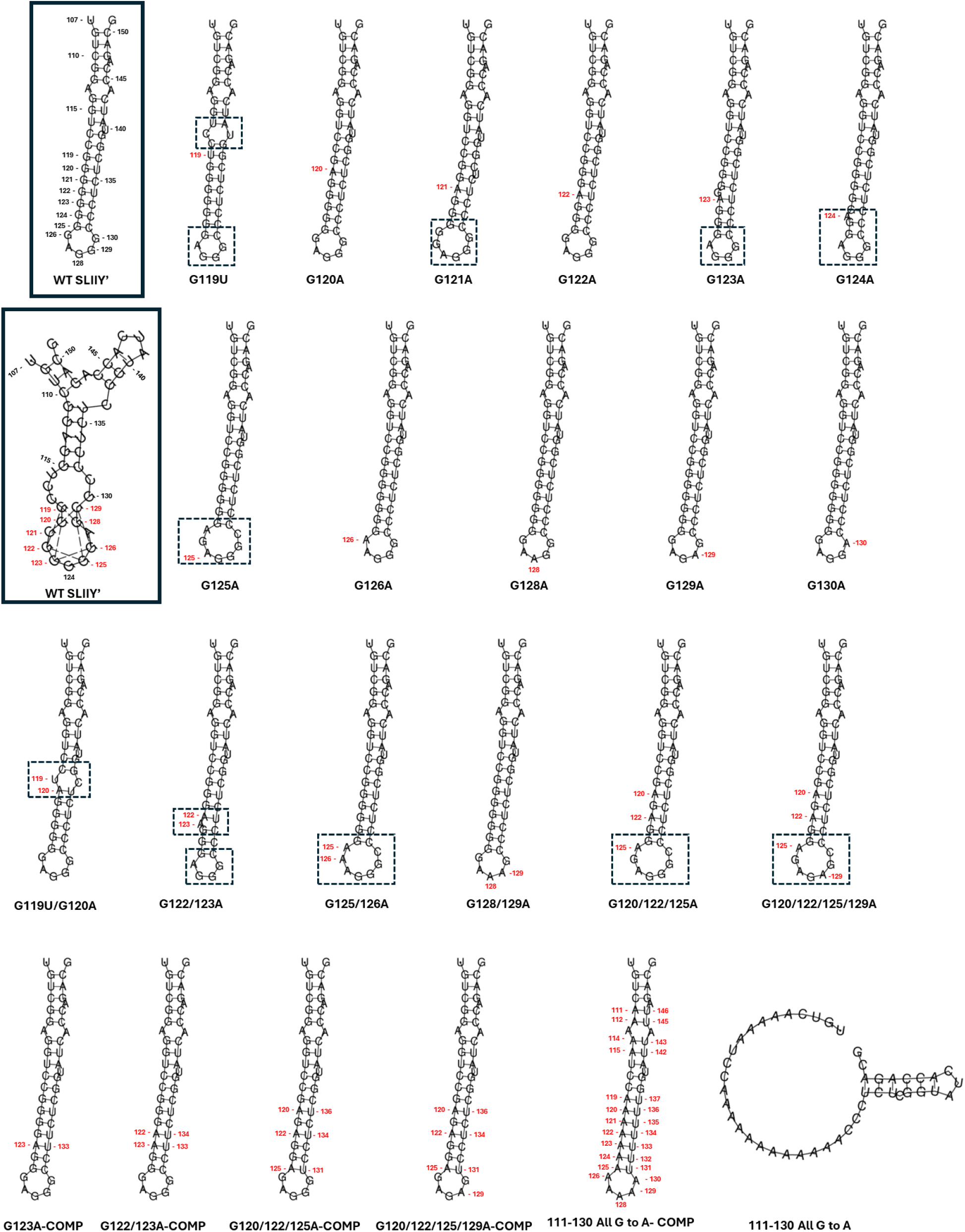
RNAfold predicted secondary structures of wild-type and mutant SL-IIy’ (nucleotides 107–151). The figure illustrates predicted folding patterns, emphasizing differences between the wild-type and mutant sequences. SL-IIy’ structures with and without G4 prediction selected are boxed. Mutated nucleotides are highlighted in red, with most mutations involving a guanine-to-adenine substitution, except for G119U, where guanine is replaced with uracil to mimic all genotypes except genotype 2. COMP denotes compensatory mutations introduced on the right side (C→U) of the hairpin to maintain the original secondary structure when mutations occur on the left side (G→A). The dashed box indicates structural changes in the hairpin following mutation compared to the wild type.

Single mutations were introduced at positions 119, 120, 121, 122, 123, 124, 125, 126, 128, 129, and 130 **(Figure 5)**. At position 119, a G→U mutation was selected because this variation is commonly found in all HCV genotypes except genotype 2, which corresponds to the JFH-1 strain used in this replicon system. The G119U mutation slightly alters the SL-IIy’ secondary structure by introducing an additional bulge and shifting one nucleotide in the bottom loop. G→A mutations at positions 121, 124, and 125 affected the loop length and hairpin size, whereas single mutations at positions 120, 122, 126, 128, 129, and 130 did not significantly impact the overall predicted SL-IIy’ structure. The double mutant G119U/G120A produced a bulge similar to that seen with G119U alone, while G122A/G123A altered loop length and stem structure. Larger structural changes were observed in G125A/G126A, G120A/G122A/G125A, and G120A/G122A/G125A/G129A, all of which extended the loop length within the stem-loop structure.

Mutations within the loop, such as G128A/G129A, did not affect the overall stem-loop conformation despite changes in the primary sequence. However, mutating all guanines to adenines from positions 111 to 130 completely disrupted the hairpin structure. To distinguish between structural and sequence-dependent effects, compensatory mutations on the right hand side of the hairpin (C→U) were introduced to stabilize hairpin formation while preserving the wild-type secondary structure.

Replicon assays were performed using wild-type, replication-deficient (ΔGDD), and successive G→A mutant constructs **(Figure 6)**. In vitro transcribed (+) strand HCV replicon RNA was electroporated into Huh7.5 cells and harvested at 4-, 24-, 48-, and 72-hours post-electroporation. To account for variations in transfection efficiency, luciferase activity was normalized to the 4-hour value of the construct with the highest luciferase signal. The 4-hour time point is critical in this assay, as this is the time when translation occurs, but there is no HCV RNA replication. The normalized luciferase output for each construct was then adjusted using the calculated ratio and plotted as a time course **(Figure 6 a-d)**. Single G→A mutants were compared to wild-type and ΔGDD and appear to replicate very similar to wild-type levels, with the exception of G129A, which has very low replication levels similar to ΔGDD. When looking at the introduction of successive G→A mutations **(Figure 6c)**, G125/126A, G128/129A, G128/129A, G120/122/125A, AND G120/122/125/129A all have significantly low **(Figure 6e)** viral replication at the 48 hour time point. The same low-level replication is observed for the compensatory mutants **(Figure 6d, 6e),** which retain hairpin secondary structure despite primary sequence mutation.

**Figure 6.**
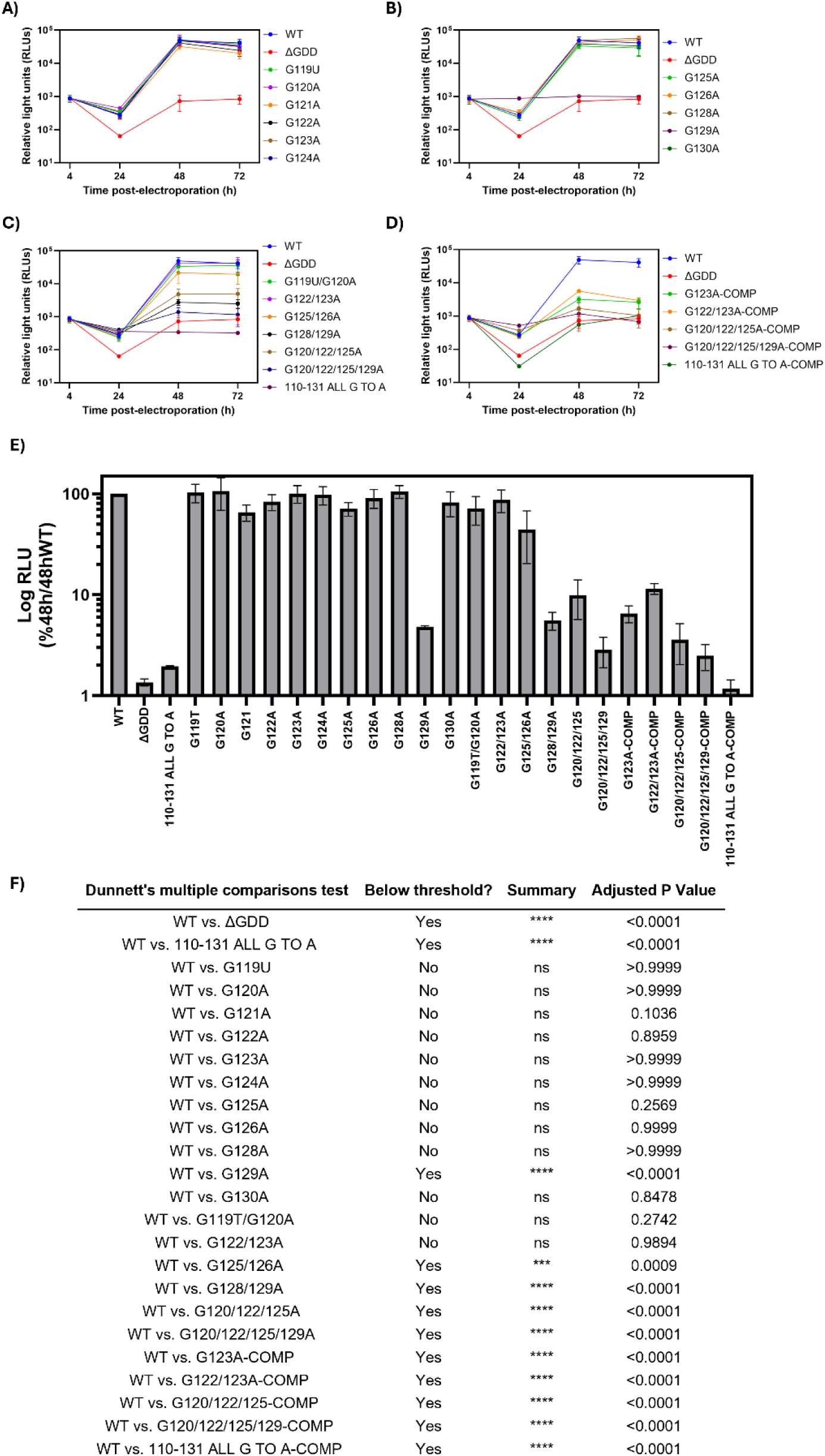
Mutations that stabilize hairpin formation or disrupt G4 formation in SL-IIy’ impair (+) RNA synthesis from the (−) strand in HCV. In vitro-transcribed wild-type or mutant HCV, replicons were transfected into Huh7.5 cells and harvested at 4-, 24-, 48-, and 72-hours post-electroporation. To account for differences in transfection efficiency over time, luciferase activity was normalized to the 4-hour value. For the 24–72 h time points, half the number of cells were plated, and equal amounts of lysate and Fluc luciferase assay buffer were incubated and measured using a luminometer. ΔGDD represents an inactive replicon with a deletion in the active site of the NS5B RNA polymerase. Time-course data are shown for **(A)** single G→A mutations, **(B)** double mutants, **(C)** triple mutants, **(D)** quadruple and compensatory mutants. **(E)** The normalized 48-hour luciferase value is presented as a percentage of wild-type activity, as the 48-hour time point best reflects RNA replication. **(F)** Statistical differences between mutant and wild-type replication at 48 hours were assessed using Dunnett’s multiple comparisons test. Mean values from three biological replicates are shown.

To support that luciferase activity can be used as an appropriate proxy for HCV RNA levels, total HCV RNA levels were quantified for each mutant **(supplemental figure S35)**. A larger variability was observed in HCV RNA levels between samples, compared to results in the luciferase assays. This can likely be attributed to sample processing or input amounts for the RT-qPCR assays. As observed in the luciferase assays, a similar decrease in RNA accumulation was observed for all replicons from 4h-24h to 24 h post-electroporation. A reduction in RNA accumulation from 4h-72h was observed for the replication dead ΔGDD construct, G129A, G128/129A, G120/122/125A, G120/122/125/129A, G122/123A-COMP, G120/122/125A-COMP, G120/122/125/129A-COMP, 110-131 All G→A COMP; which is also consistent with the luciferase output data presented in **Figure 6**.

## Discussion

Recent studies have identified guanine quadruplexes (G4s) in several flaviviruses, including DENV, ZIKV, WNV, and HCV, suggesting that these non-canonical RNA structures may influence viral replication, translation, and genome stability(38–42). To investigate the role of these G4s, many studies utilize bioinformatics to identify potential G4-forming sequences and biophysical methods such as circular dichroism spectroscopy and NMR to evaluate their formation *in vitr*o. Cell culture assays are used to elucidate the role of these G4s during viral replication. However, these studies largely employ small-molecule compounds (e.g., TMPyP4, PhenDC3, BRACO-19) that stabilize these structures(43). The interpretation of these cell culture results can be challenging, as G4s are found not only in the viral genomes but also throughout the host cellular genome in promoter regions and telomeric ends(44). As such, it is difficult to determine whether the antiviral inhibitory effects are due to the stabilization of G4s in the cellular genome or the viral genome, as these G4s can often regulate the transcription initiation of many host genes that are required for viral replication.

In this study, we built upon prior observations of Friebe and Bartenschlager (19), in which the authors identified that the last 157 nucleotides at the 3′ end of the HCV (−) strand is the minimal sequence required for efficient RNA replication. Based on the secondary structure of the negative strand 3’UTR determined by footprinting experiments by Smith et al. (18) and mutational analysis by Friebe and Bartenschlager (2009), the SL-I’, SLIIz’ are required for RNA replication, while SLIIy’ significantly contributes to RNA replication. Based on HCV-replicon assays, the authors conclude that for SLIIy’ the primary RNA sequence is more important than the stem structure for efficient replication (19). Upon bioinformatic analysis of the HCV genome for G4s, Jaubert et al. (2018) (11) identified a potential G-quadruplex forming sequence in the negative strand of HCV located in the SLIIy’ (nucleotides 110-131) (11). The authors performed extensive biophysical analysis using CD-spectroscopy and NMR to show that nucleotides 110-131 form a G4 *in vitro*. To assess the biological significance of this G4 structure, they performed in vitro RNA synthesis assays. They demonstrated that G4 formation inhibits RNA-dependent RNA polymerase (RdRp) activity, directly interfering with viral RNA synthesis. Additionally, they evaluated the effect of Phen-DC3, a well-characterized G4 ligand, and found that it effectively inhibits HCV replication in cell culture under conditions without cytotoxicity. The limitation of the work performed in cell culture is that the Phen-DC3 small molecule non-specifically binds to any G4s intracellular and viral and can confound the results interpreted.

In the current study, we expanded on the findings observed in the previously described papers (11, 18, 19) and utilized bioinformatics to verify that across 339 sequences obtained from genotypes 1-7, a highly conserved G4 is present at nucleotide positions 110-131 **(Figure 1 and Table 1)**. We performed extensive biophysical characterization using CD-spectroscopy **(Table 1 and Figure 2)**, which shows that the wild-type negative-strand G4 on SLIIy’ (nucleotides 110-131nt) forms a parallel-stranded G4, consistent with the findings from Jaubert et al(11). To systematically investigate the impact of G4 disruption, we chose to introduce successive guanine-to-adenine mutations as a model for destabilizing G4 structures(22). However, across all seven genotypes, the HCV positive-strand 5’UTR is the most conserved region in the genome, with approximately 5-10% sequence divergence among genotypes(45). The 5′UTR is highly structured and can be divided into four structural domains (I, II, III, and IV), which include miR-122 binding sites (nucleotides 21-23 and 38-40) and the IRES (nucleotides 39-371)(46). Due to the essential role of the positive-strand 5’UTR in replication, translation, and interaction with crucial host factors, introducing mutations into this region can negatively impact translation and genome stability.

Given that disrupting IRES function would confound our ability to specifically study the effects of 5’UTR mutations on RNA replication, we opted to utilize a well-established HCV replicon system developed by Friebe and Bartenschlager (19). This system provides a strategic advantage, as it enables the introduction of mutations in the 5’UTR without affecting translation efficiency **(Supplemental S36)**. In this replicon construct **(Figure 4, Supplemental S2)**, the 5’UTR is separated from the rest of the non-structural proteins and the 3’UTR, creating a poly-cistronic reporter system where the Fluc reporter gene is placed under the control of the poliovirus IRES, ensuring that reporter translation remains unaffected by modifications to the HCV 5’UTR. Meanwhile, the translation of the HCV structural and non-structural proteins is driven by the Encephalomyocarditis virus IRES. Therefore, under these conditions, any mutations introduced into the 5’UTR of the HCV positive strand will selectively impact RNA replication initiation from the 3’UTR of the HCV negative strand, allowing us to isolate the specific effects of these modifications on viral RNA replication.

We demonstrated through CD-spectroscopy **(Table 1**, **Figure 2)**, SAXS **(Figure 3**, **Table 2)**, bioinformatic analysis using G4-hunter **(Table 1)** and structural prediction algorithm RNA-fold **(Table 1**, **Figure 5)** that successive guanine to adenine mutations disrupts quartet formation of the G4s and there are observable structural changes. The G4 hunter algorithm, where a score of >1 indicates that a G4 is likely to form in the sequence, predicted that all the single mutant, double mutant and the G120/122/129A or G122/125/129A triple mutants could form G4s. However, based on the CD spectroscopy data and SAXS data, this is not the case. Single point mutations of G129A and G119T appear to disrupt quartet formation as the peak at ∼260 nm for CD spectroscopy with peak ellipticity values of 5.8 mdeg and 6.8 mdeg, respectively when compared to the wild-type of 14.3 mdeg. Successive double, triple, and quadruple mutations severely impacted the peak amplitude at ∼260 nm **(Table 1)**. Moreover, the RNA fold algorithm computationally predicted that a G4 would be able to form with mutations G120A, G123A, and G129A; however, based on the CD spectra and SAXS profiles compared to wild-type 110-131, they appear to be different.

Based on the prediction algorithms and biophysical data, a select number of successive G→A mutants were selected to test in the HCV replicon system **(Figure 4**, **Figure 5)**. A comprehensive panel of single mutants were chosen to elucidate the impact of G→A mutation at a single-nucleotide level. Double and triple G→A mutants were chosen based on their biophysical results and computational prediction on their ability to disrupt G4 structure. Moreover, four additional mutants were chosen where a G→A mutation was introduced, but a complementary C→U mutation was introduced to support the formation of a stem-loop structure instead of a guanine quadruplex (labelled COMP). A negative control where all guanines were mutated to adenines from nucleotides 110-131 was also included in the replicon assays.

The replicon assay results **(Figure 6)** indicated that G129A replicated at significantly lower levels compared to the wild-type HCV replicon. The CD-spectroscopy and SAXS data, also support that G129A mutations disrupt guanine quadruplex mutations, indicating that a single nucleotide mutation in the loop sequence at position 129 is sufficient to disrupt quadruplex formation. The RNA-fold predicted G4 secondary structure **(Figure 5)** suggests that G129A stabilizes at least two possible quartets, which may explain why mutations at this position are poorly tolerated. Double mutant G128/129A also has the lowest replication amongst the tested replicon double mutants, providing further evidence that the loop sequence disruption is likely an important location for G4 formation due to its accessibility and flexibility. Although G119U appears to have the G4 quartets disrupted from biophysical characterization of the HCV 110-131 oligo, G4hunter and RNA fold show that a G4 is still likely to form. Moreover, this mutant was still able to replicate efficiently in the luciferase assays, indicating that there is some discrepancy between the biophysical data and the computational and cell culture data. The G→A quadruple mutant G120/122/125/129A also has extremely low levels of replication, likely not only due to G4 quartet disruption as indicated by CD-spectroscopy and a distinct SAXS envelope with 3 domains but also increased loop size compared to wild-type **(Figure 5)**. Stabilization of the hairpin structure through G123A-COMP and G122/123A-COMP indicates an approximate 90% reduction in viral replication, indicating that the hairpin structure still contributes to RNA replication but is likely responsible for a minority of the importance. While successive mutations with 3 or more G→A mutations, which have been compensated to stabilize the hairpin secondary structure, negatively impact viral replication significantly. These findings indicate that mutations in the SLIIy’ are not well tolerated, and stabilization of the stem-loop structure does not restore viral replication to wild-type levels. As observed by Friebe and Bartenschlager, the primary sequence in this region is likely more important than the hairpin secondary structure (19). We have shown that mutations that do not disrupt quadruplex formation (i.e. all single mutants and double mutants that do not include G129A), as demonstrated by CD Spectroscopy and SAXS, do not significantly impact viral replication. While disruption of the G4 or stabilization of the hairpin confirmation negatively impacts viral replication.

Mechanistically, a study performed by Belachew et al. (9) identified that the HCV NS3 helicase plays a crucial role in viral replication, not only by unwinding duplex RNA during replication but also by resolving stable secondary structures such as G-quadruplexes (G4s). NS3 is a member of the DExH helicase family, which belongs to superfamily 2, exhibiting unwinding activity on both RNA and DNA substrates by translocating in the 3’ to 5’ direction (47, 48). Like other helicases in this family, such as DHX36 and DHX9, NS3 plays a crucial role in resolving stable secondary structures like G4RNAs, thereby removing obstacles for transcription and translation machinery (49–51). A key finding in the study conducted by Bleachew et al. (2022) is that the NEGG4 sequence, which represents the G4-forming sequence found in SLIIy’ of the HCV negative strand, forms a stable two-tetrad G4 structure that must be unfolded for productive viral replication (9). *In vitro* studies indicated that NS3 exhibited significant G4-unfolding activity on NEGG4, and the presence of G4-stabilizing ligands, such as Phen-DC3, significantly inhibited NS3’s unfolding activity, further supporting the idea that NS3 actively resolves G4s as part of its function in viral RNA synthesis. Given that NS3 is likely the only helicase present in the membranous web, the site of HCV replication, it may be uniquely adapted to regulate G4 structures in the HCV genome, ensuring that these stable elements do not impede RNA synthesis (52, 53)

From our experiments, combined with previous findings from (9, 11, 18, 19), we propose that the G4 structure within SLIIy’ plays a critical role in recruiting the HCV NS3 helicase, which is essential for unwinding secondary RNA structures. This unwinding process is necessary to facilitate access for the viral RNA-dependent RNA polymerase, NS5B, ensuring efficient genome replication. Given that SLIIy’ may adopt an alternative, more energetically favourable G4 structure rather than the predicted stem-loop conformation, it is possible that the formation of this G4 serves as a regulatory switch that modulates helicase recruitment and replication efficiency. In the absence of a G4 structure, NS3 recruitment may be impaired, leading to inefficient RNA unwinding and reduced polymerase activity, ultimately hindering viral replication. These findings highlight a previously unrecognized functional role of G4 structures in the HCV replication cycle as a regulator for (+) strand RNA synthesis.

Future studies could further investigate the mechanistic interactions between NS3, NS5B, and the SLIIy’ G4, as well as assess whether this mechanism is conserved in other G4-forming regions of HCV. Additionally, these observations and established HCV replicon systems could be leveraged to study G4s identified in the 5′ UTR of the positive strand and the 3′ UTR of the negative strand in other flaviviruses. Since HCV shares key replication mechanisms with other positive-sense single-stranded RNA (+ssRNA) flaviviruses, such as dengue virus (DENV), Zika virus (ZIKV), and West Nile virus (WNV), it serves as a valuable model for investigating how G4 structures regulate viral replication, translation, and genome stability across the Flaviviridae family. Insights gained from these studies could help uncover conserved G4-associated regulatory mechanisms in related flaviviruses, potentially revealing new targets for broad-spectrum RNA-targeting antivirals.

## Supplementary Materials

The following supporting information can be found along with the published article. S1: Sequences of all samples used for nucleotide 110-131 sequence alignment; S2: wild-type plasmid sequence for pFK341-Sp-PI-luc-EI-core-3′JFH1 replicon system provided by Dr. Ralf Bartenschlager; Figure S3: Size exclusion chromatograms of all oligos used in this study; S4-S34: SAXS – raw data, Guinier plot, Kratky plot, P(r) plot, and low-resolution SAXS envelopes for all oligos used in this study; S35: HCV RNA accumulation for wild-type and mutant replicons; S36: Replicon data for all replication dead ΔGDD wild-type and mutant constructs (110-131nt).

## Author Contributions

C.S.C and T.R.P.: conceptualization of the study and preparation of the article; S.D. conceptualized the study, designed and completed the experiments, analyzed the data, and prepared the article; S.T and M.D.B.: assisted with the SAXS analysis and preparation of the article; A.C. and M.S.H assisted with sample processing for cell culture experiments. G.v.M. and J.A.C. helped with the study design and preparation of the article.

## Acknowledgements

We extend our gratitude to Prof. Ralf Bartenschlager for generously sharing the pFK341-Sp-PI-luc-EI-core-3′JFH1 replicon system and to Paul Rothhaar for helpful discussions on the luciferase assays. We are also grateful to Prof. Stephan Urban for providing the Huh7.5 cell line. We appreciate Deepak Patel from Dr. Alexei Savchenko’s lab for the use of the Akta Pure system for SEC purification of all oligonucleotides used in this study. Additionally, we are grateful to Marylin Rheault and Manolya Sag from Dr. Selena Sagan’s lab for their valuable discussions that significantly enhanced the optimization of our HCV replicon assays and in-house RT-qPCR. We are also thankful to Michael D’souza in Dr. Trushar Patel’s lab for helping setup the instrument for CD spectroscopy. We thank DIAMOND Light Source and B21 staff members for this help with SAXS data collection.

## Institutional Review Board Statement

Ethics approval was obtained from the University of Calgary for the use of cell lines by Conjoint Health Research Ethics Board, CHREB (Ethics ID: REB20-0933).

## Data Availability Statement

The data that support the findings of this study will be shared on reasonable request to the corresponding author.

